# Tracking the outbreak. An optimized delimiting survey strategy for *Xylella fastidiosa*

**DOI:** 10.1101/2020.03.05.978668

**Authors:** E. Lázaro, M. Sesé, A. López-Quílez, D. Conesa, V. Dalmau, A. Ferrer-Matoses, A. Vicent

## Abstract

- Current legislation enforces the implementation of intensive surveillance programs for quarantine plant pathogens. After an outbreak, surveys are implemented to delimit the geographic extent of the pathogen and execute disease control. The feasibility of control programs is highly dependent on budget availability, thus it is necessary to target and optimize surveillance strategies.
- A sequential adaptive delimiting survey involving a three-phase and a two-phase design with increasing spatial resolution was developed and implemented for the *Xylella fastidiosa* outbreak in Alicante, Spain. Inspection and sampling intensities were optimized using simulation-based methods and results were validated using Bayesian spatial models.
- This strategy made it possible to sequence inspection and sampling considering different spatial resolutions, and to adapt the inspection and sampling intensity according to the information obtained in the previous, coarser, spatial resolution.
- The proposed strategy was able to delimit efficiently the extent of *Xf* improving efficiency of the current in terms of survey efforts. From a methodological perspective, our approach provides new insights of alternative delimiting designs and new reference sampling intensity values.

## 1 Introduction

Regulation (EU) 2016/2031 (EU, 2016) and Implementing Regulation (EU) 2019/2072 (EU, 2019b) define the list of quarantine plant pests and pathogens for which the adoption of measures is necessary to prevent their introduction into and spread within the European Union (EU). As indicated by these Regulations, Member States should take all necessary phytosanitary measures to eradicate quarantine pests and pathogens, when found to be present in their territories. In those cases where a quarantine plant pest or pathogen cannot be longer eradicated, containment measures may apply. Nevertheless, the efficacy of eradication or containment measures strongly relies on a precise delimitation of the infested area. Accurate diagnostic protocols exist for regulated plant pathogens and are routinely used by plant health authorities when tracking quarantine outbreaks (Petter & Suffert, 2010). However, surveillance and sampling methodologies are far from optimal representing a serious bottleneck for the effective implementation of control measures.

*Xylella fastidiosa* (*Xf*) is a xylem-inhabiting phytopathogenic bacterium (Wells *et al*., 1987) which can cause different diseases in a wide range of cultivated, ornamental, and forest plant species (EFSA, 2018a; Saponari *et al*., 2019). Due to its potential economic, environmental and social impacts, *Xf* is included in the list of priority quarantine pests and pathogens for the EU (EU, 2019a). Member States shall carry out specific surveys including a sufficiently high number of visual examinations, sampling and testing. However, *Xf* infections can manifest in different ways, from latent asymptomatic forms to a quick plant dieback due to complex interactions among the host, pathogen, and environment, which sometimes make visual detection difficult (Purcell *et al*., 1999; Loconsole *et al*., 2016).

The bacterium is a genetically diverse species grouped into six subspecies, although *Xf* subsp. *fastidiosa, Xf* subsp. *pauca, Xf* subsp. *multiplex*, and *Xf* subsp. *sandyi* are the four most frequently reported (Schaad *et al*., 2004; Denancé *et al*., 2017). Nevertheless, only the subspecies *fastidiosa* and *multiplex* are recognized by the Committee on the Taxonomy of Plant Pathogenic Bacteria of the International Society of Plant Pathology (ISPP) (Bull *et al*., 2012). *Xf* subsp. *fastidiosa* has been found in grapevines, citrus, coffee, and almond; *Xf* subsp. *pauca* has been found in citrus and coffee; *Xf* subsp. *multiplex* has been found in almond, peach, plum, oak, blueberry, pecan, etc.; and *Xf* subsp. *sandy* has been found in oleander.

The species *Xf* develops in the vascular system of the plants and it is naturally transmitted by xylem sap-feeding insects, which spread the pathogen to relatively short distances (Almeida *et al*., 2014). Long-distance spread is usually associated with human activities that involve moving infected plant hosts or vectors (Nunney *et al*., 2014; Morente *et al*., 2018). Originally, the geographic distribution of the bacterium was restricted to the Americas, where it was endemic (EFSA, 2019). However, its presence was recently confirmed in Iran (Amanifar *et al*., 2014) and Taiwan (Su *et al*., 2016), and since 2013 it is officially present in the European Union (EU), specifically in Italy (2013), France (2015), Spain (2016), and Portugal (2018) (see EC, 2019, for further details).

The current situation of *Xf* in Europe is raising major concerns that are giving rise to a phytosanitary emergency. Two of the most relevant crops in the Mediterranean areas of the EU, olives and almonds, are severely affected by *Xf* diseases. Grapevines were also found to be affected in the Balearic Islands, Spain, and other major crops such as citrus are at risk. Furthermore, the insect vector *Philaenus spumarius* (meadow spittlebug) is widespread in affected regions in Spain and Italy as well as other areas of the EU territory (Saponari *et al*., 2014; Cornara *et al*., 2017; EFSA, 2018b).

After first being detected in the EU, additional emergency measures were enforced for *Xf* to prevent further spread under Decision 2015/789/EU (EU, 2015b) (hereinafter referred to as “the Decision”) and its subsequent amendments 2015/2417/EU, 2016/764/EU, 2017/764/EU, 2018/927/EU and 2018/1511/EU (EU, 2015a, 2016, 2017, 2018b,a). Among other actions, the Decision establishes the implementation of two different surveillance actions, detection (Art. 3) and delimiting (Art. 6) surveys, depending on the pathogen status in the area. Detection surveys are aimed at detecting the pest and ensuring the status of “pest-free area”. After the first detection of *Xf* in an area, delimiting surveys are then conducted to demarcate the boundaries and geographic extent of the pathogen (Art. 4). Surveillance activities are pivotal to enhance *Xf* control (i.e., eradication or containment) and to understand the epidemiology and dynamics of the disease (EFSA, 2016). Consequently, the development of innovative and advanced methods for delimiting or detection has been prioritized by the European authorities (see EFSA, 2016, for further details).

Considering the regulatory framework for *Xf* in the EU as a starting point, the present work focuses on developing an alternative strategy for delimiting surveys in the demarcated area of the province of Alicante, Spain, as a case study. The outbreak in Alicante was first reported in June 2017 and since then delimiting survey activities have been conducted, covering a demarcated area of about 140,000 ha. Almond (*Prunus dulcis*) is the most affected plant species, although others such as *Rosmarinus officinalis, Polygala myrtifolia, Helichrysum italicum* and various shrub plants have also been detected, but representing a minor proportion. So far, only *Xf* subsp. *multiplex* has been detected in the demarcated area (Giampetruzzi *et al*., 2018). With circa 2,000 orchards and 51,000 trees already destroyed (GVA, 2019), this outbreak represent one of the largest plant disease eradication campaigns ever attempted in Europe. Nevertheless, the feasibility of this control program is seriously compromised by the low efficiency of the current surveillance strategy for *Xf* enforced by Decision 2015/789/EU.

Delimiting surveys are fundamental to assess the feasibility of disease management plans and target control tactics (Stanaway, 2011; Potts *et al*., 2013; Tobin *et al*., 2013; Hauser *et al*., 2016). Implemented after an initial detection, delimitation implies inspection (i.e., visual examination of plants) and sampling actions and its effectiveness is highly dependent on budget availability, which ends up limiting resources for monitoring and control. Thus, there is a need to design delimiting surveys which minimize the costs of inspection while maintaining an acceptable level of risk of overlooking a positive (Hauser *et al*., 2016).

During the last two decades innovative survey methods have been developed taking into consideration optimization-based tools with the aim of improving their efficiency (Epanchin-Niell & Liebhold, 2015; Büyüktahtakin & Haight, 2018). However, the number of studies published on this topic is scarce and they rarely consider the particular case of delimiting strategies (Hauser *et al*., 2016; Moore & McCarthy, 2016; Yemshanov *et al*., 2017a,b).

As has been suggested by Pacifici *et al*. (2016), recent advances in surveillance methodologies focus not only on improving the sampling design but also on developing more sophisticated data analyses to overcome deficiencies in the survey design. In relation to the sampling methodology, there are several strategies such as sequential sampling (Chaudhuri & Stenger, 2005), stratified sampling (Edwards *et al*., 2005), or adaptive sampling (Brown *et al*., 2013) which can increase the information content and provide a more efficient estimation when disease distribution is spatially correlated (Pacifici *et al*., 2016).

On the other hand, the use of complex statistical models can overcome deficiencies in the survey design and data collection. For instance, spatial autocorrelation and other factors associated with imperfect survey techniques such as observer error, sampling gaps or missing data can now be included in the modeling process given the developments in statistical and computing methods (Latimer *et al*., 2006; Banerjee *et al*., 2014; Martínez-Minaya *et al*., 2018). Bayesian statistics allow all these data particularities to be introduced into the model structure, and more especially Bayesian hierarchical models have been proven to be a good option to deal with spatial correlation and other dependence structures (Latimer *et al*., 2006; Banerjee *et al*., 2014).

In our study, an alternative delimiting survey strategy was developed for the particular case of *Xf* in Alicante. This strategy was defined by a sequential adaptive scheme which combines different spatial resolution grid sizes in different survey phases. The sequential adaptive survey begins with a full inspection of all grid cells and a simple random sampling. This is followed by a second phase of additional inspection and sampling at a higher spatial resolution, but only in those cells that were found to be infested in the first phase of the survey. This sequential adaptive strategy was implemented considering a two-phase and a three-phase design. For each phase, optimum inspection and sampling intensities were estimated using simulation-based optimization methods to ensure the efficacy was comparable to that of the current strategy (Decision 2015/789/EU).

An additional objective was also considered in our study, namely, to determine the influence of the survey design on the estimates of the spatial distribution of *Xf* incidence in the demarcated area in Alicante. Incidence (i.e., the proportion of plants positive for *Xf*) is a magnitude commonly used in plant pathology to characterize the disease status or to evaluate the efficacy of a control program (Madden *et al*., 2007). A Bayesian hierarchical spatial model was used here to infer this metric using data from the official surveys and simulated data obtained considering the outputs of the sequential adaptive strategy indicated above. A comparison among the inferential outputs obtained with the different survey designs was carried out quantifying differences in the stability of the estimates using several discrepancy measures.

## 2 Material and methods

### 2.1 Database

In compliance with the current legal provisions, after the detection of *Xf* in 2017 the competent plant health authority carried out delimiting surveys in the demarcated area in Alicante. These surveys were aimed at updating the current extent and boundaries of the infested area, assessing pathogen incidence, and implementing eradication measures as enforced by Decision 2015/789/EU. In the present study, data from the 2018 delimiting survey campaign (hereinafter referred to as the “2018 official survey”) were used, The dataset used included a total of 8,142 samples from individual plants covering a total of 83,300 ha.. Samples were analyzed for the presence of *Xf* in official laboratories following the EPPO standard diagnostic protocol (EPPO, 2019). Additional information related to sampling date, plant species, presence of symptoms (i.e., symptomatic vs. asymptomatic), and GPS coordinates (in WGS84 reference system) were also collected.

Altogether 124 species were sampled in the 2018 official survey with *Olea europaea, Prunus dulcis, Ficus carica, Rosmarinus officinalis*, and *Vitis* spp. accounting for 62.06% of the total sample size (Table 1). Plants expressing *Xf* -like symptoms were preferentially sampled, representing 85% of the total. In the laboratory analysis, 3.37% (227/6957) of the symptomatic samples were found to be positive for *Xf*, while only 0.84% (10/1185) were positive in the case of the asymptomatic samples. Altogether, 237 samples were positive for *Xf* with 221 of them belonging to *Prunus dulcis*, while the rest corresponded to *R. officinalis* (1), *R. alaternus* (1), *P. armeniaca* (1), *P. myrtifolia* (5), *Phagnalon saxatile* (3), *Helycrhysum italicum* (3), *Calicotome spinosa* (1), and *Scabiosa atropurpurea* (1).

**Table 1:**
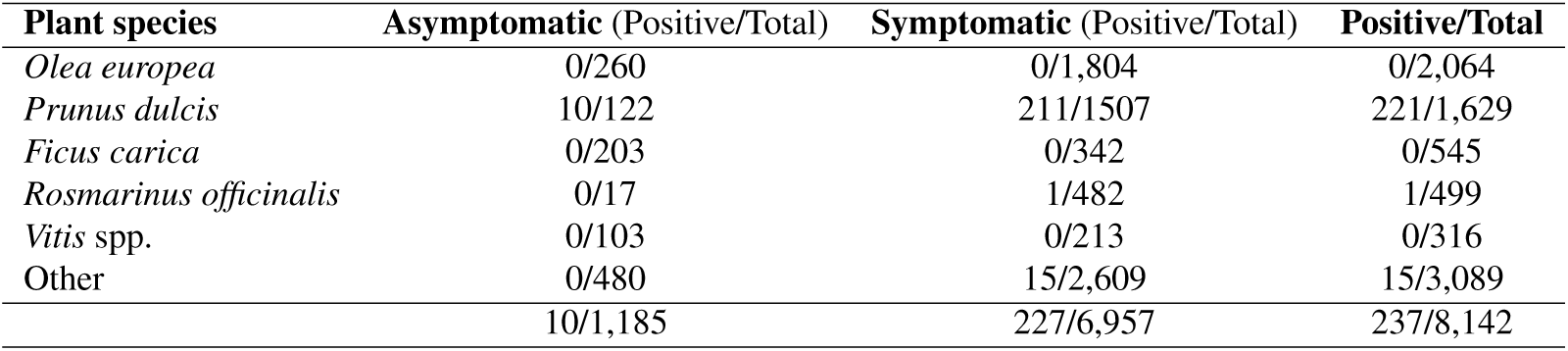
Absolute frequency distribution of the presence of symptoms and positive detection of *Xylella fastidiosa* in the dataset of the 2018 official survey in Alicante categorized by plant species.

| Plant species | Asymptomatic (Positive/Total) | Symptomatic (Positive/Total) | Positive/Total |
| --- | --- | --- | --- |
| *Olea europaea* | 0/260 | 0/1,804 | 0/2,064 |
| *Prunus dulcis* | 10/122 | 211/1507 | 221/1,629 |
| *Ficus carica* | 0/203 | 0/342 | 0/545 |
| *Rosmarinus officinalis* | 0/17 | 1/482 | 1/499 |
| *Vitis* spp. | 0/103 | 0/213 | 0/316 |
| Other | 0/480 | 15/2,609 | 15/3,089 |
|  | 10/1,185 | 227/6,957 | 237/8,142 |

### 2.2 Evaluation of delimiting strategies

#### Current delimiting strategy

In compliance with Art. 4 of the Decision, under an eradication situation, delimiting surveys are aimed at establishing a “demarcated area” with an infested zone (i.e., infected zone, in the Decision) surrounded by a buffer zone. The infected zone must include all plants infected by *Xf* and all plants in the vicinity liable to be infected and/or showing symptoms within a radius of 100 m. The buffer zone is delimited by considering a 5 km radius around the infected zone. However, this distance may be reduced to no less than 1 km under certain conditions or increased up to 10 km under a containment situation (see EU, 2015b, for further details). Additionally, the buffer zone must be intensively monitored and maintained under disease-free (i.e., pest-free, in the Decision) conditions. This intensive monitoring is performed at different spatial resolutions: i) 100 *x* 100 m grid cells (0.01 km^2^) in the 1^*st*^ km radius, and ii) on 1 *x* 1 km grid cells (1 km^2^) in the rest of the buffer zone. Monitoring involves visual inspections of plants, which should be sampled and tested for *Xf* preferably when observing symptoms.

In line with the legal provisions in force, in 2018 the delimiting surveys in Alicante resulted in 71 infected zones and an aggregated buffer zone split into 656 cells of 1 km^2^ and 17,700 cells of 0.01 km^2^. The extent of the buffer zone covered a total of 833 cells of 1 km^2^, with 177 of them being re-split into 0.01 km^2^ cells. The grid layout of the demarcated area is shown in Fig. 1(a) and the distribution of samples can be seen in Fig. 1(b).

**Fig. 1:**
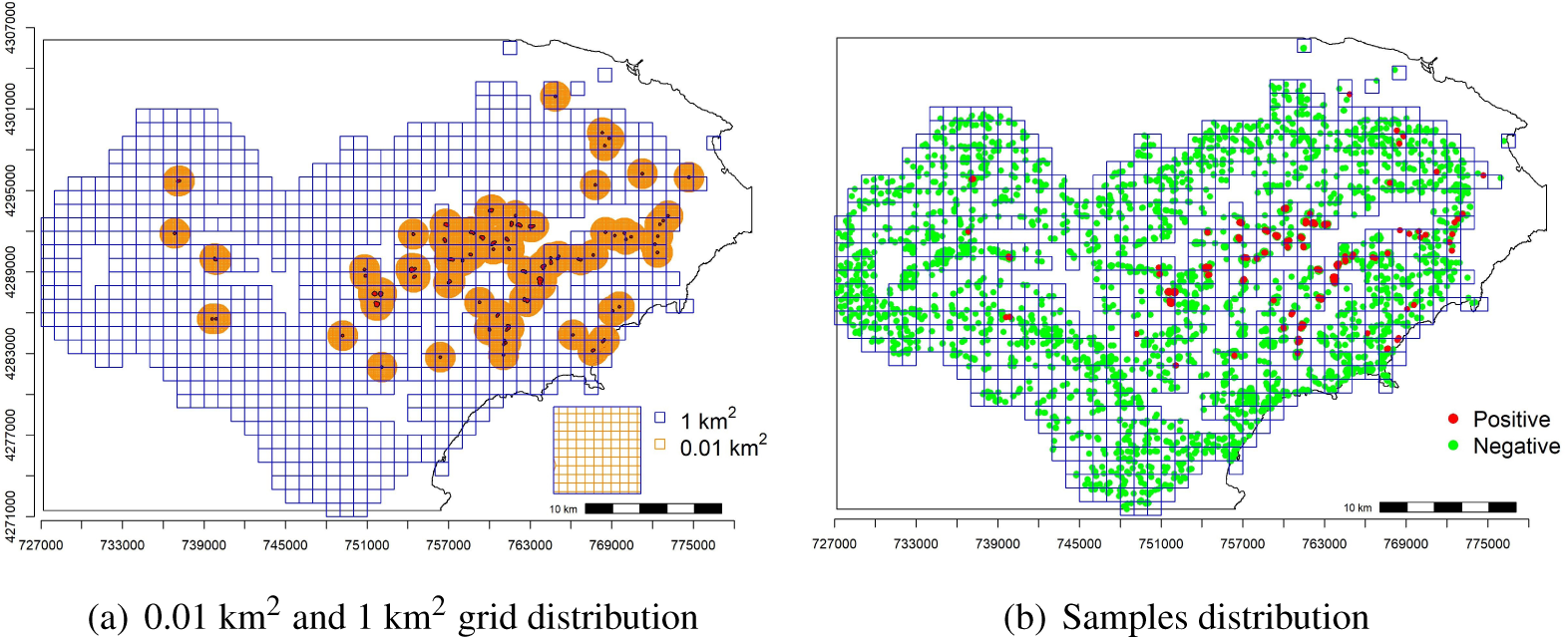
Distribution of (a) 0.01 km^2^ and 1 km^2^ grid, and (b) samples giving positive and negative for *Xylella fastidiosa* in the demarcated area in Alicante, Spain, in 2018.

In terms of the sampling description, Table 2 provides information about the number of sampled cells, sampling intensity (number of samples/cell), and additional information categorized by grid resolution. Note that a grid of 500 m x 500 m (0.25 km^2^) was also included in the description as being later considered in one sequential adaptive design (see next section). Considering the 1 km^2^ grid, the percentage of cells sampled reached 100% with an overall rate of positives for *Xf* of 8.53%, and median and maximum sampling intensity values of 5 and 109 samples/cell, respectively. For the 0.25 km^2^ and 0.01 km^2^ grids, the percentage of sampled cells was 49.33% and 4.01%, respectively. The overall percentage of positives for *Xf* was 2.67% for the 0.25 km^2^ grid and 0.19% for the 0.01 km^2^ grid. The median sampling intensity was 1 sample/cell in both grid resolutions, with a maximum of 101 for the 0.25 km^2^ grid and 59 samples/cell for the 0.01 km^2^ grid.

**Table 2:** Sampling description in the demarcated area in Alicante, Spain, for *Xylella fastidiosa* in 2018 categorized by different grid resolutions (*j* = {1, 0.25, 0.01} km^2^). ***C*** _*j*_ denotes the number of cells covering the demarcated area depending on the grid resolution (*j*); *C*_*j,s*_ is the number of cells per grid (*j*) in which at least one sample was taken; and *C*_*j*,+_ is the number of cells per grid (*j*) in which at least one sample was detected as positive for *X. fastidiosa*. Sampling intensity (samples/cell) is described by the median and the maximum values.

| Grid resolution<br>(km<sup>2</sup>) | Number of cells |  |  | Sampling intensity |  |
| --- | --- | --- | --- | --- | --- |
| | $C_j$ | $C_{j,s}$ | $C_{j,+}$ | Median | Maximum |
| $j = 1$ | 833 | 833 | 71 | 5 | 109 |
| $j = 0.25$ | 3,332 | 1,644 | 89 | 1 | 101 |
| $j = 0.01$ | 83,100 | 3,340 | 161 | 1 | 59 |

#### Sequential adaptive strategy

An alternative delimiting strategy was proposed under the premises of improving the efficiency and maintaining the efficacy of the current strategy established by the Decision. To improve the efficiency, inspection intensity (i.e., number of cells inspected for each grid resolution) and sampling intensity (i.e., number of samples taken in each cell) were optimized using a sequential adaptive scheme which combines 1 km^2^, 0.25 km^2^, and 0.01 km^2^ grid resolutions in different phases of the survey. Assuming an initial survey resolution of 1 km^2^, the sequential approach allows the phases of the survey to be scheduled in different time frames. Additionally, the adaptive approach allows the inspection and sampling intensities to be tailored for each phase depending on the results obtained in the previous phase at a coarser spatial resolution.

Our proposal considers inspection and sampling of all 1 km^2^ cells in the demarcated area and inspection and sampling are performed at a finer spatial resolution only in those cells in which *Xf* was detected. This adaptive sequence was implemented by a three-phase and a two-phase design, which differed in the sequence of grid resolutions. The three-phase design included increasing grid resolutions of 1 km^2^, 0.25 km^2^, and 0.01 km^2^, whereas the two-phase design only included 1 km^2^ and 0.01 km^2^. Fig. 2 illustrates the phase sequences of the two designs with an example. This scheme was applied to the 2018 official survey database, which was considered the reference to optimize inspection and sampling intensities.

**Fig. 2:**
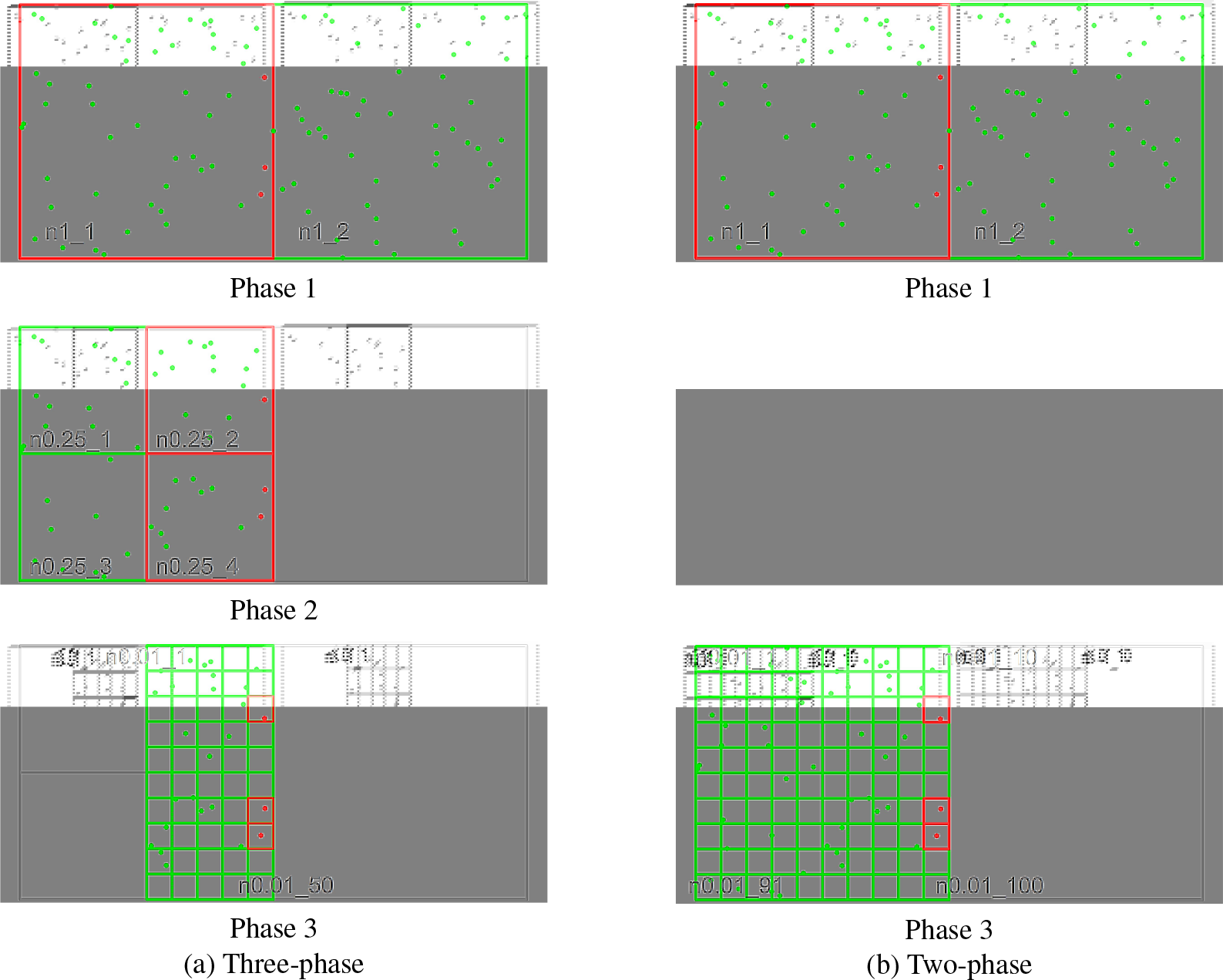
Example of the sequential adaptive strategy for a) the three phase and b) the two phase designs for 1 km^2^ (row 1), 0.25 km^2^ (row 2), and 0.01 km^2^ (row 3) grid resolutions. Red and green denote positive and negative samples (dots) or cells (squares), respectively, for *Xylella fastidiosa*. Gray cells represent those not inspected or sampled for a specific grid resolution. *n* _*j*_ for *j* = {1, 0.25, 0.01} denotes sampling intensity (samples/cell) for the different grids.

**Fig. 3:**
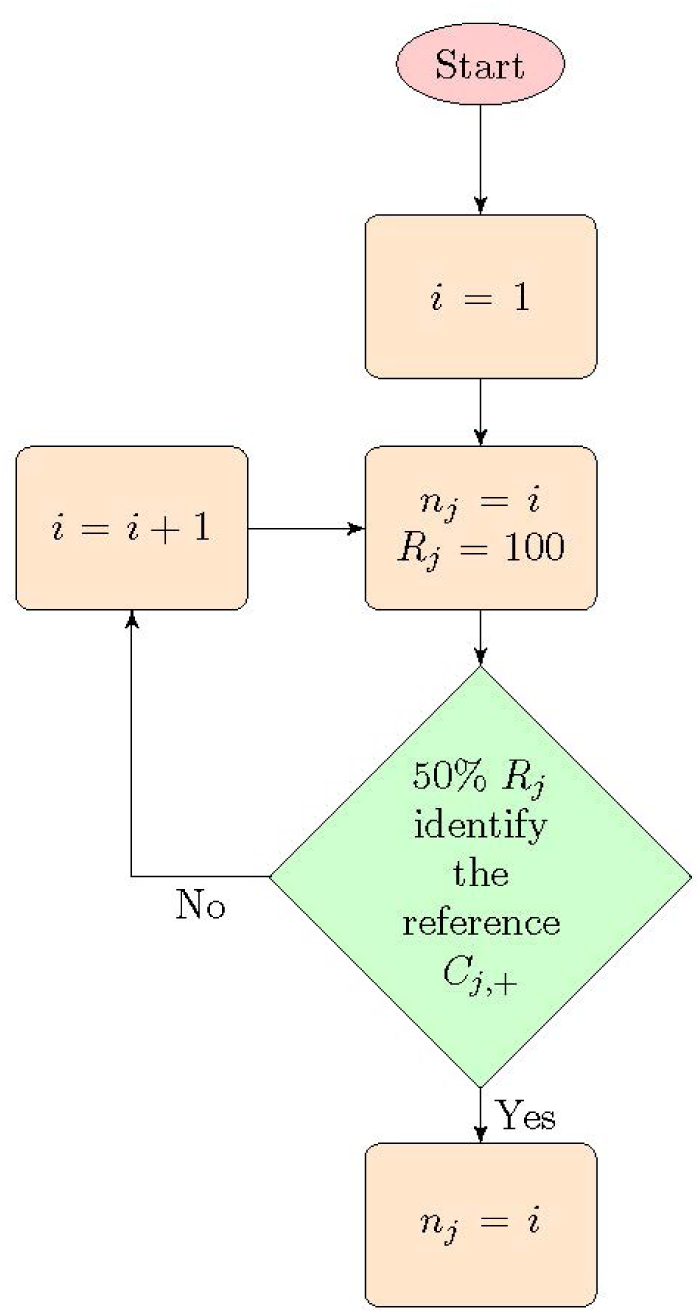
Optimization algorithm to estimate optimum sampling intensity *n* _*j*_ for *j* = {1, 0.25, 0.01} in the three-phase design and *j* = {1, 0.01} for the two-phase design. *R*_*j*_ and *C*_*j*,+_ denote the number of replicates and the number of positive cells for *Xylella fastidiosa* in each *j* grid resolution.

*Phase 1*. For all 1 km^2^ cells:

*Step 1*. Initialize *n*_1_ as 1, where *n*_1_ denotes the optimum sampling intensity for the 1 km^2^ grid resolution. This value is the threshold sampling intensity value.
*Step 1b*. Increment *n*_1_ by 1.
*Step 2*. Run 100 replicates (*R*_1_ = 100) of a random sampling under the restriction imposed by the value *n*_1_.
*Step 3*. Stopping rule: if 50% of the replicates (*R*_1_) have identified all the positive cells for *Xf* in the 1 km^2^ grid resolution (*C*_1,+_), *n*_1_ is the optimum sampling intensity. If not, go back to *step 1b* and continue the sequence.

*Phase 2*. Only for 1 km^2^ cells identified as positive for *Xf* in *phase 1*. (Phase 2 is not considered in the two-phase design):

*Step 1*. Initialize *n*_0.25_ as 1, where *n*_0.25_ denotes the optimum sampling intensity for the 0.25 km^2^ grid resolution. This value is the threshold sampling intensity value.
*Step 1b*. Increment *n*_0.25_ by 1.
*Step 2*. Run 100 replicates (*R*_0.25_ = 100) of a random sampling under the restriction imposed by the value *n*_0.25_.
*Step 3*. Stopping rule: if 50% of the replicates (*R*_0.25_) have identified all the positive cells for *Xf* in the 0.25 km^2^ grid resolution (*C*_0.25,+_), *n*_0.25_ is the optimum sampling intensity. If not, go back to *step 1b* and continue the sequence.

*Phase 3*. Only for 0.25 km^2^ cells identified as positive for *Xf* in *phase 2* (three-phase design) or 1 km^2^ cells identified as positive in *phase 1* (two-phase design):

*Step 1*. Initialize *n*_0.01_ as 1, where *n*_0.01_ denotes the optimum sampling intensity for the 0.01 km^2^ grid resolution. This value is the threshold sampling intensity value.
*Step 1b*. Increment *n*_0.01_ by 1.
*Step 2*. Run 100 replicates (*R*_0.01_ = 100) of a random sampling under the restriction imposed by the value *n*_0.01_.
*Step 3*. Stopping rule: if 50% of the replicates (*R*_0.01_) have identified all the positive cells for *Xf* in the 0.01 km^2^ grid resolution (*C*_0.01,+_), *n*_0.01_ is the optimum sampling intensity. If not, go back to *step 1b* and continue the sequence.

According to the survey scheme illustrated in Fig. 2 and the outputs of the optimization algorithm (*n*_1_, *n*_0.25_, *n*_0.01_), the survey efforts (i.e., the total number of samples) for the three-phase and two-phase designs were calculated as:

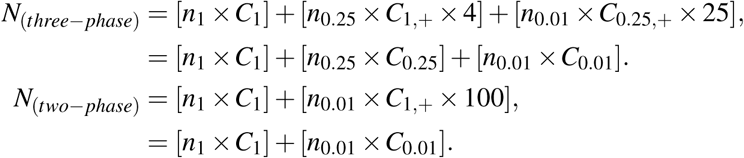

where *C*_1_, *C*_0.25_, *C*_0.01_ are the total number of cells to be inspected in the 1 km^2^, 0.25 km^2^, and 0.01 km^2^ grid resolutions; *C*_1,+_ and *C*_0.25,+_ are the number of cells identified as positive for *Xf* in the 1 km^2^ and 0.25 km^2^ grid resolutions; and *n*_1_, *n*_0.25_, *n*_0.01_ are the optimum sampling intensities (samples/cell) calculated by the algorithm for the 1 km^2^, 0.25 km^2^, and 0.01 km^2^ grid resolutions. Note that the equivalence of grid cells at increasing spatial resolutions for the three-phase design are 1 km^2^ → 0.25 km^2^ = 4 and 0.25 km^2^ → 0.01 km^2^ = 25. Likewise, the equivalence of grid cells for the two-phase design is 1 km^2^ → 0.01 km^2^ = 100.

Delimiting strategies were evaluated by comparing the three-phase and two-phase sequential adaptive designs with the current strategy based on the estimation of inspection and sampling intensities. Inspection and sampling intensity outputs were combined to provide an overall evaluation of survey efforts.

### 2.3 Modeling the distribution of *Xf* incidence

#### Bayesian hierarchical spatial model

Based on previous work (Cendoya *et al*., unpublished), the spatial variation of *Xf* incidence was modeled here by means of a Bayesian hierarchical spatial model. This methodology allows the introduction of more stochasticity in the model by means of spatial structured effects (Latimer *et al*., 2006) and other non observed sources of variability as random effects, thereby improving the accuracy of the estimates, uncertainty quantification, and predictive power. The inference process was addressed under the integrated nested Laplace approximation (INLA) proposed by Rue *et al*. (2009) and implemented through the `R-INLA` package (see Martínez-Minaya *et al*., 2018, for further details on methodological assumptions to apply the INLA approach).

This analysis was performed using the data of the 2018 official survey, georeferenced to the regular lattice of 1 km^2^ cells used for the surveillance of the demarcated area. Given the nature of the database, an extension of a generalized linear model (GLM) (Nelder & Wedderburn, 1972) was proposed to define a generic model as follows:

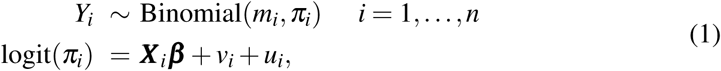

in which the number of samples positive for *Xf* in each grid cell (*Y*_*i*_) was considered as the response variable and as following a Binomial distribution, *Y*_*i*_∼ *Binomial*(*m*_*i*_; *π*_*i*_), with *π*_*i*_ and *m*_*i*_ denoting the probability of a sample being positive for *Xf* and the total number of samples in cell *i*, respectively. The linear predictor was defined by a vector of covariates and its corresponding vector of coefficients, ***X*** _*i*_ and ***β***, and by a spatial and an independent random effect associated to each cell *i, v*_*i*_, and *u*_*i*_, respectively. This generic model is usually known as the Besag, York and Molli é model (Besag *et al*., 1991) and it makes it possible to take into account similarities among neighboring cells and also to quantify intra-cell ability to be positive for *Xf* (unstructured random effects).

In particular, random effects were defined as a Gaussian distribution with mean 0 and precision *τ*_*u*_,

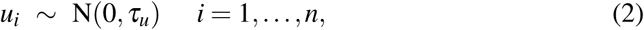

while spatial dependence was modeled considering an intrinsic conditional autoregressive structure (ICAR) (Besag, 1974). That is, each grid cell follows a conditional Gaussian distribution with mean equal to the average of the neighboring cells (structured random effects) and a precision proportional to the number of them:

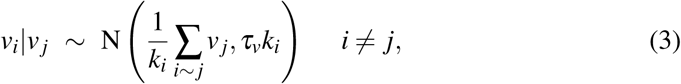

where *j* ∼ *i* denotes *i* and *j* neighboring cells, *τ*_*v*_ is the precision of the spatial random effect, and *k*_*i*_ is the number of neighbors of the corresponding cell *i*. The neighborhood criterion was established at a maximum distance of 2.5 km among all the cells to ensure that all of them had at least one neighbor. Note that this spatial structure definition accomplishes Markovian properties and it can be considered a latent Gaussian Markov random field (GMRF) (Rue *et al*., 2009, 2017), thus making the INLA implementation feasible. Prior to including the spatial effect in the model, Moran’s I test was used to check for the existence of spatial autocorrelation in the *Xf* incidence distribution (Dormann *et al*., 2007).

Spatially gridded climatic data (30” arc min resolution) from the demarcated area in Alicante were acquired from the WorldClim v.2 database (Fick & Hijmans, 2017), which contains mean monthly temperature and precipitation values for the period 1970-2000. Three bioclimatic covariates were included as fixed effects in the generic model: annual mean temperature (°C) (coded as *bio*1), temperature annual range (°C) (coded as *bio*7), and precipitation of the wettest month (mm) (coded as *bio*13). The coordinate system WGS84 was used in all spatially gridded datasets with raster package for `R` software (Hijmans, 2019).

The model formulation was completed with the elicitation of a prior distribution for the parameters and hyperparameters. Following the hierarchical structure, after defining the model likelihood (see equation (1)), priors for the parameters are specified together with the definition of random effects (see equation (2) and (3)), and lastly the prior distribution of the hyperparameters (hyperpriors) are specified as follows:

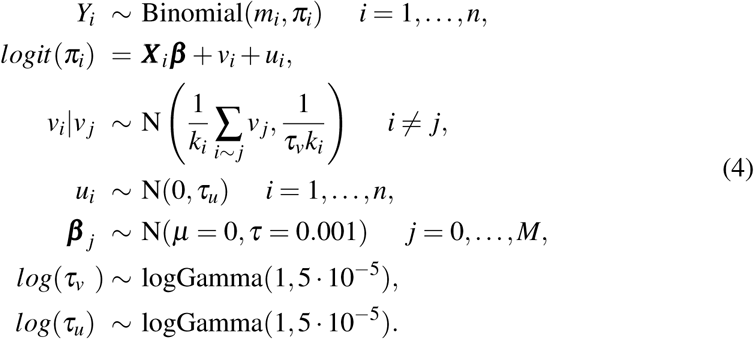

Note that a non-informative scenario was considered to set prior specification with normal distributions centered at zero and a small precision for regression coefficients and log-gamma priors (a default option of INLA) with a wide mean and variance for precisions of the spatial and independent random effects.

With the aim of selecting the more parsimonious model and with the best explanatory and predictive abilities, all possible model component combinations, 2^*k*^ (with *k* denoting the number of components of the linear predictor including random effects), were assessed in terms of goodness of fit, complexity, and predictive ability. For this purpose, two selection model criteria were used: the Watanabe-Akaike information criteria (WAIC) (Watanabe, 2010; Gelman *et al*., 2014), and the logarithmic conditional predictive ordinate (LCPO). While WAIC evaluates goodness of fit and model complexity, LCPO addresses predictive ability. The models with the lowest values of WAIC and LCPO were chosen.

#### Effect of sampling intensity on *Xf* incidence estimates

To evaluate the influence of sampling intensity on the estimates of *Xf*, different data subsets were built from the reference database of the 2018 official survey using the regular lattice of 1 km^2^ cells. Four data subsets were obtained by limiting the maximum sampling intensity to: i) the optimum sampling intensity calculated with the optimization algorithm for the 1 km^2^ grid resolution (Section 2.2); ii) the third quartile of the sampling intensity in the reference database; and iii) two arbitrary sampling intensities in between those values (Section 3.2).

The data subset generation process consisted in running a simple random sampling for each 1 km^2^ cell in which the sampling intensity was set according to the thresholds indicated above. Each data subset was replicated 100 times (*R* = 100) to ensure the inclusion of a wide range of sampling configurations. Subsequently, the data subsets were used to run the Bayesian hierarchical spatial model obtained previously for the reference dataset and selected based upon WAIC and LCPO criteria.

The 100 replicas (*R* = 100) were considered in each inferential process, so 100 independent posterior inferences were obtained and averaged to carry out the contrast. The comparison was focused on assessing the stability of: i) the marginal posterior distribution of the fixed parameters (regression coefficients) and hyperparameters (spatial and independent effects standard deviation), ii) the marginal posterior distribution of the incidence, iii) the standard deviation associated to the posterior distribution of the incidence, and iv) the marginal posterior distribution of the spatial and the independent effects. Note that spatial and independent random effects were characterized by their corresponding standard deviation: 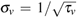 and 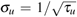.

The stability of the marginal posterior distributions of the fixed parameters and hyperparameters was evaluated by means of the following discrepancy measures:

- **Bias**: Difference between the average of the posterior sample means of the replicas and the posterior sample mean of the reference inference process, 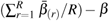, where *R* is the number of replicas (*R* = 100 in our study),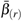 is the mean of the posterior marginal corresponding to the replica *r*, and *β* is the mean of the posterior marginal corresponding to the reference model.
- **Standard error (SE)**: Square root of the average of the posterior variances of the replicas,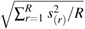, where 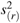 is the sample variance of the posterior sample for replica *r*.
- **Standard deviation (SD)**: Standard deviation of the set 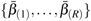 that includes the mean of the posterior marginal of the regression coefficients (or hyperparameters) of all replicas.

Note that 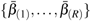represents the vector of means of the marginal posterior distributions for each the regression parameter (denoted as *β*) corresponding to the replica *r, r* = 1, …, 100, of the inferential process. For assessing hyperparameter discrepancy measures we proceeded analogously but focused on 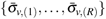and 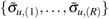.

On the other hand, the stability of the derived quantities (posterior distribution of the incidence and its corresponding standard deviation, spatial and independent effects) was assessed by means of the bias, computed as the difference between the average of the posterior sample means (for the derived quantities) of the replicas and the posterior sample mean of the reference inference process. Note that the reference inference process was defined by the model fitted for the reference dataset and selected based upon WAIC and LCPO criteria.

## 3 Results

### 3.1 Evaluation of delimiting strategies

#### Inspection intensity

Table 3 contains the graphical and numerical description of the inspection intensity for the two sequential adaptive designs (three-phase and two-phase) and the current strategy. The inspection intensity of the current strategy in 1 km^2^ (656 cells) was lower than in both sequential adaptive designs (833 cells), given that the first kilometer of the buffer zone must be inspected directly at a resolution of 0.01 km^2^. In contrast, the inspection intensity at a grid resolution of 0.01 km^2^ was lower for both alternative designs (2225 (three-phase) and 7100 (two-phase) cells). Specifically, the three-phase design had the lowest inspection intensity value (2225 cells) due to the integration of the intermediate grid resolution of 0.25 km^2^, which substantially reduced the inspection intensity.

**Table 3:**
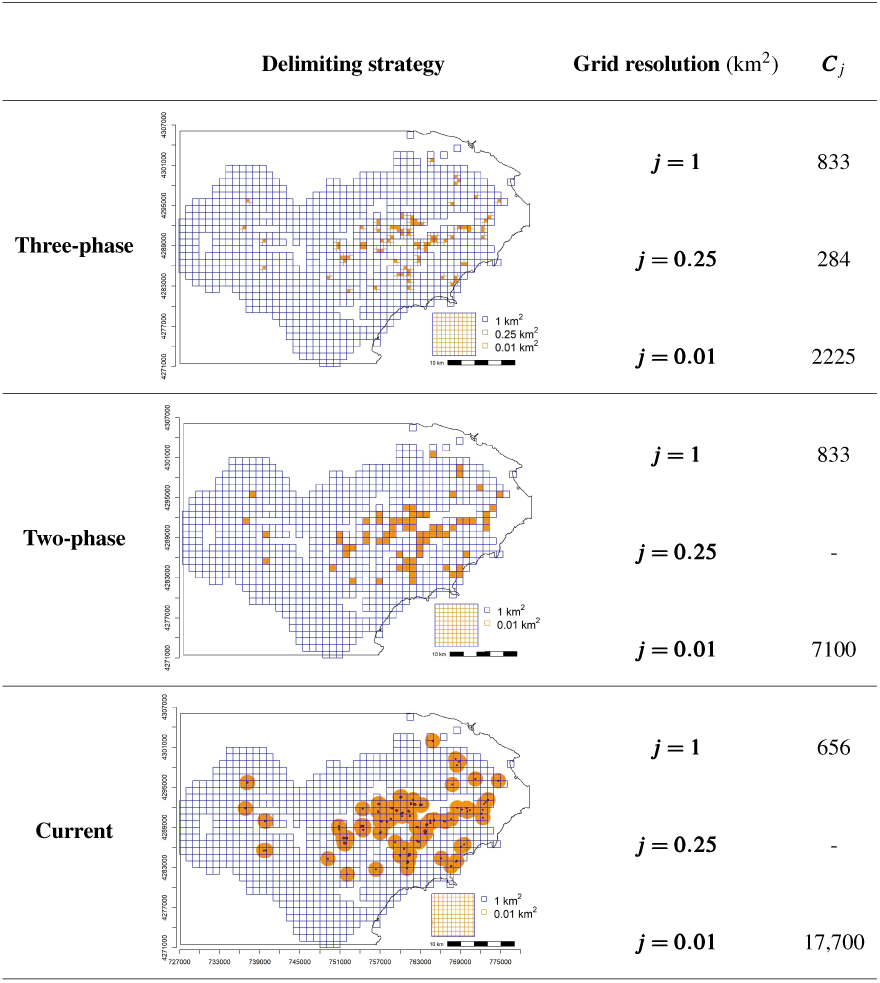
Inspection intensity for the three-phase, two-phase, and current delimiting strategies in the demarcated area for *Xylella fastidiosa* in Alicante, Spain. ***C*** _*j*_ denotes the inspection intensity as the number of cells to be inspected for each grid resolution *j* = {1, 0.25, 0.01} km^2^.

| | Delimiting strategy | Grid resolution ($\text{km}^2$) | $C_j$ |
| --- | --- | --- | --- |
| **Three-phase** | | $j = 1$ | 833 |
| | | $j = 0.25$ | 284 |
| | | $j = 0.01$ | 2225 |
| **Two-phase** | | $j = 1$ | 833 |
| | | $j = 0.25$ | - |
| | | $j = 0.01$ | 7100 |
| **Current** | | $j = 1$ | 656 |
| | | $j = 0.25$ | - |
| | | $j = 0.01$ | 17,700 |

#### Sampling intensity

Table 4 shows the optimum sampling intensities obtained for the three-phase and two-phase sequential adaptive designs under the established condition of *R*_*j*_ = 50% and with other less restrictive conditions (*R*_*j*_ = 25%, *R*_*j*_ = 15% and *R*_*j*_ = 5%). Note that, given the sequential adaptive nature of the strategy, the condition *R*_*j*_ established for a particular phase (*j*) affected the following one. 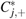 denotes the number of positive cells for *Xf* that our algorithm identified for the specific optimum sampling intensity (Table 4) summarized by the median value of the replicates.

**Table 4:**
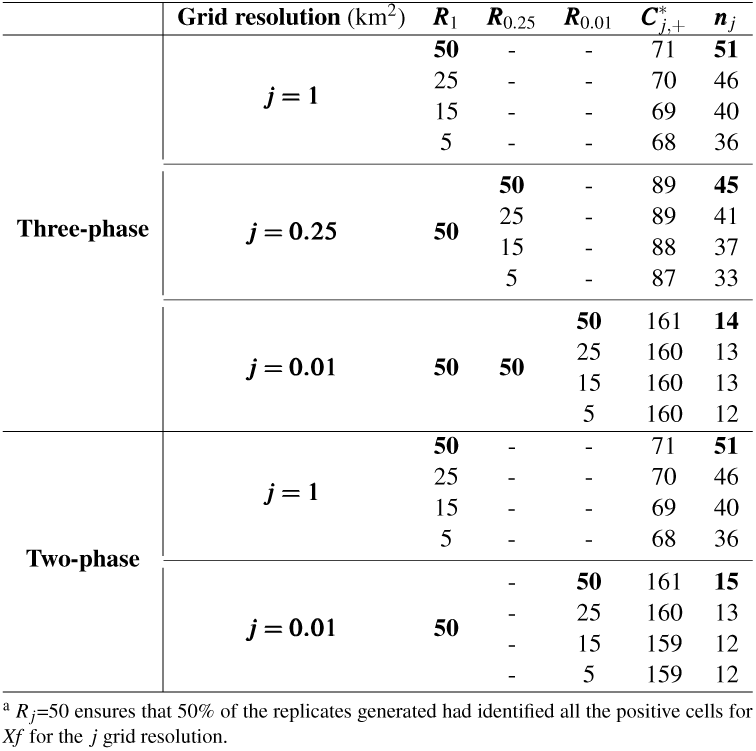
Optimum sampling intensity (*n* _*j*_) calculated for each grid resolution *j* = {1, 0.25, 0.01} in the three-phase and two-phase sequential adaptive designs applied to the demarcated area for *Xylella fastidiosa* in Alicante, Spain. *R*_*j*_ ^a^ denotes the condition established in the optimization algorithm for each grid resolution and 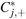 is the number of positive cells for *Xf* identified by the algorithm under the constraint imposed.

| | Grid resolution (km<sup>2</sup>) | $R_1$ | $R_{0.25}$ | $R_{0.01}$ | $C_{j,+}^*$ | $n_j$ |
| --- | --- | --- | --- | --- | --- | --- |
| Three-phase | $j = 1$ | **50** | - | - | 71 | **51** |
|  |  | 25 | - | - | 70 | 46 |
|  |  | 15 | - | - | 69 | 40 |
|  |  | 5 | - | - | 68 | 36 |
| | $j = 0.25$ | **50** | **50** | - | 89 | **45** |
|  |  |  | 25 | - | 89 | 41 |
|  |  |  | 15 | - | 88 | 37 |
|  |  |  | 5 | - | 87 | 33 |
| | $j = 0.01$ | **50** | **50** | **50** | 161 | **14** |
|  |  |  |  | 25 | 160 | 13 |
|  |  |  |  | 15 | 160 | 13 |
|  |  |  |  | 5 | 160 | 12 |
| Two-phase | $j = 1$ | **50** | - | - | 71 | **51** |
|  |  | 25 | - | - | 70 | 46 |
|  |  | 15 | - | - | 69 | 40 |
|  |  | 5 | - | - | 68 | 36 |
| | $j = 0.01$ | **50** | - | **50** | 161 | **15** |
|  |  |  | - | 25 | 160 | 13 |
|  |  |  | - | 15 | 159 | 12 |
|  |  |  | - | 5 | 159 | 12 |

<sup>a</sup> $R_j=50$ ensures that 50% of the replicates generated had identified all the positive cells for *Xf* for the $j$ grid resolution.

As shown in Table 4, for the three-phase design optimum sampling intensity was calculated in 51, 45 and 14 samples/cell for *j* = {1, 0.25, 0.01} km^2^ grid resolutions. These optimum values ensured the detection of all the positive cells for *Xf* given that 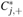 values matched *C*_*j*,+_ (Table 2). For 1 and 0.25 km^2^ grid resolutions, differences in sampling intensity varied a maximum of 15 (51-36) and 12 (45-33) samples/cell among the different conditions established by the algorithm, respectively, whereas differences in 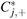 were no more than 3 (71-68) and 2 (89-87) positive cells each. This showed that a relatively small number of cells were sampled more intensively than the others. Sampling intensity decreased with higher grid resolution, although differences between 1 and 0.25 km^2^ grid resolutions were not greater than 6 samples/cell.

For the two-phase design, as shown in Table 4, the optimum sampling intensity was calculated in 51 and 15 samples/cell for *j* = {1, 0.01} km^2^ grid resolutions. All these optimum values ensured the detection of all positive cells for *Xf* (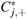 values that matched *C*_*j*,+_ displayed in Table 2). For 0.01 km^2^ differences in sampling intensity values varied a maximum of 3 (15-12) samples/cell among the different conditions established by the algorithm, and differences in 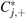 were no more than 2 (161-159) positive cells. Sampling intensity decreased with higher grid resolution and the difference between 1 and 0.01 km^2^ corresponded to a value of 36 (51-15) samples/cell for the most restrictive condition.

#### Survey effort

An overall assessment of the survey effort of the proposed sequential adaptive designs (three-phase and two-phase) and the current delimiting survey strategy is provided in Table 5.

**Table 5:** Overall assessment of the three-phase and two-phase sequential adaptive designs and the current delimiting survey strategy in the demarcated area for *Xylella fastidiosa* in Alicante, Spain. For each grid resolution *j* with *j* = {1, 0.25, 0.01*}, C*_*j*_ denotes the inspection intensity (number of cells to be inspected), *n* _*j*_ is the sampling intensity (number samples/cell), and *N*_*j*_ = *C*_*j*_ × *n* _*j*_ is the sampling effort as the total number of samples to be taken. ∑ _*j*_ *N*_*j*_ denotes total survey effort.

| | Grid resolution (km<sup>2</sup>) | $C_j$ | $n_j$ | $N_j$ |
| --- | --- | --- | --- | --- |
| **Three-phase** | $j = 1$ | 833 | 51 | 42,483 |
| | $j = 0.25$ | 284 | 45 | 12,780 |
| | $j = 0.01$ | 2,225 | 14 | 31,150 |
|  |  |  |  | **86,413** |
| **Two-phase** | $j = 1$ | 833 | 51 | 42,483 |
| | $j = 0.01$ | 7,100 | 15 | 106,500 |
|  |  |  |  | **148,983** |
| **Current** | $j = 1$ | 656 | 51 | 33,456 |
| | $j = 0.01$ | 17,700 | 15 | 265,500 |
|  |  |  |  | **298,956** |

Both sequential adaptive designs improved efficiency in terms of inspection and sampling intensities for the 0.01 km^2^ grid resolution. This result strongly influenced the overall number of samples to be taken (i.e., survey effort), which were estimated as 86,413, 148,983 and 298,956 for the three-phase, two-phase and current delimiting strategies, respectively.

Inspection and sampling intensities were lower for the three-phase design in comparison to the two-phase (2225 vs. 7100 cells, and 31,150 vs. 106,500 samples) due to the integration of an intermediate grid resolution (0.25 km^2^) which substantially reduced the overall survey effort (86,413 vs. 148,983). On the other hand, the inspection effort of the current strategy in the 1 km^2^ grid resolution was lower than that of the sequential adaptive designs given that the 1 km radius of the buffer zone is surveyed directly at a resolution of 0.01 km^2^. However, it was far greater in the 0.01 km^2^ grid resolution, the overall survey effort increasing to 298,956 samples.

### 3.2 Modeling the distribution of *Xf* incidence

#### Model selection

The following model was selected based on WAIC and LCPO criteria (see Supplementary material Table S1 for WAIC and LCPO values) among all 64 possible combinations for the 2018 official survey database:

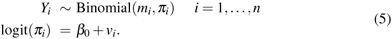

The model with the lowest WAIC value was the one which included the covariates *bio*1, *bio*7, and the spatial effect (Supplementary material Table S1). The lowest LCPO value corresponded to the model with *bio*7, *bio*13 and the spatial effect (Supplementary material Table S1). Nevertheless, models ranked first based on WAIC and LCPO presented similar scores (less than one unit of difference), so the most parsimonious model including only the spatial effect was selected. Table 6 shows a numeric descriptive of the marginal posterior parameters and hyperparameter distributions for the selected model. The full model is also described to illustrate why climatic variables were not finally included. Mean values of the posterior distributions of the climatic covariates were very close to zero and their corresponding probabilities of being greater than zero were around 0.50, thus having low explanatory capacity for the distribution of *Xf* incidence.

**Table 6:** Marginal posterior distributions of parameters and hyperparameters for the model of *Xylella fastidiosa* incidence distribution in the demarcated area in Alicante, Spain, with mean, standard deviation (**SD**), median (***Q***_0.5_), and 95% credible interval (**95% CI**). The full model including climatic covariates is also indicated.

|  | Mean | SD | Q<sub>0.5</sub> | 95% CI | P(·) > 0 |
| --- | --- | --- | --- | --- | --- |
| $\beta_0 + v$ | | | | | |
| $\beta_0$ | -5.829 | 0.325 | -5.807 | [-6.526,-5.253] | 0 |
| $\sigma_v$ | 1.762 | 0.202 | 1.748 | [1.401,2.194] | - |
| $\beta_0 + bio1 + bio7 + bio13 + v$ | | | | | |
| $\beta_0$ | -7.273 | 13.897 | -7.205 | [-34.859,19.906] | 0.298 |
| *bio1* | 0.060 | 0.312 | 0.055 | [-0.540,0.687] | 0.570 |
| *bio7* | 0.040 | 0.343 | 0.040 | [-0.634,0.717] | 0.545 |
| *bio13* | -0.008 | 0.100 | -0.005 | [-0.214,0.181] | 0.477 |
| $\sigma_v$ | 1.798 | 0.208 | 1.791 | [1.653,2.228] | - |

Fig. 4 shows the mean posterior distribution of the spatial effect, the mean posterior distribution of the incidence (0-1), and its corresponding standard deviation for the model of *Xf* incidence distribution in the demarcated area in Alicante, Spain. As indicated before, the mean posterior distribution of the spatial effect was determined by the spatial dependence structure defined by a neighborhood relation of a distance of 2.5 km among cells. It ranged from −1.780 to 4.371, with positive values indicating higher *Xf* incidence estimates. As can be observed, the central and eastern parts of the demarcated area included the cells with the highest values for the spatial effect, corresponding to those cells where a higher proportion of positives was detected.

**Fig. 4:**
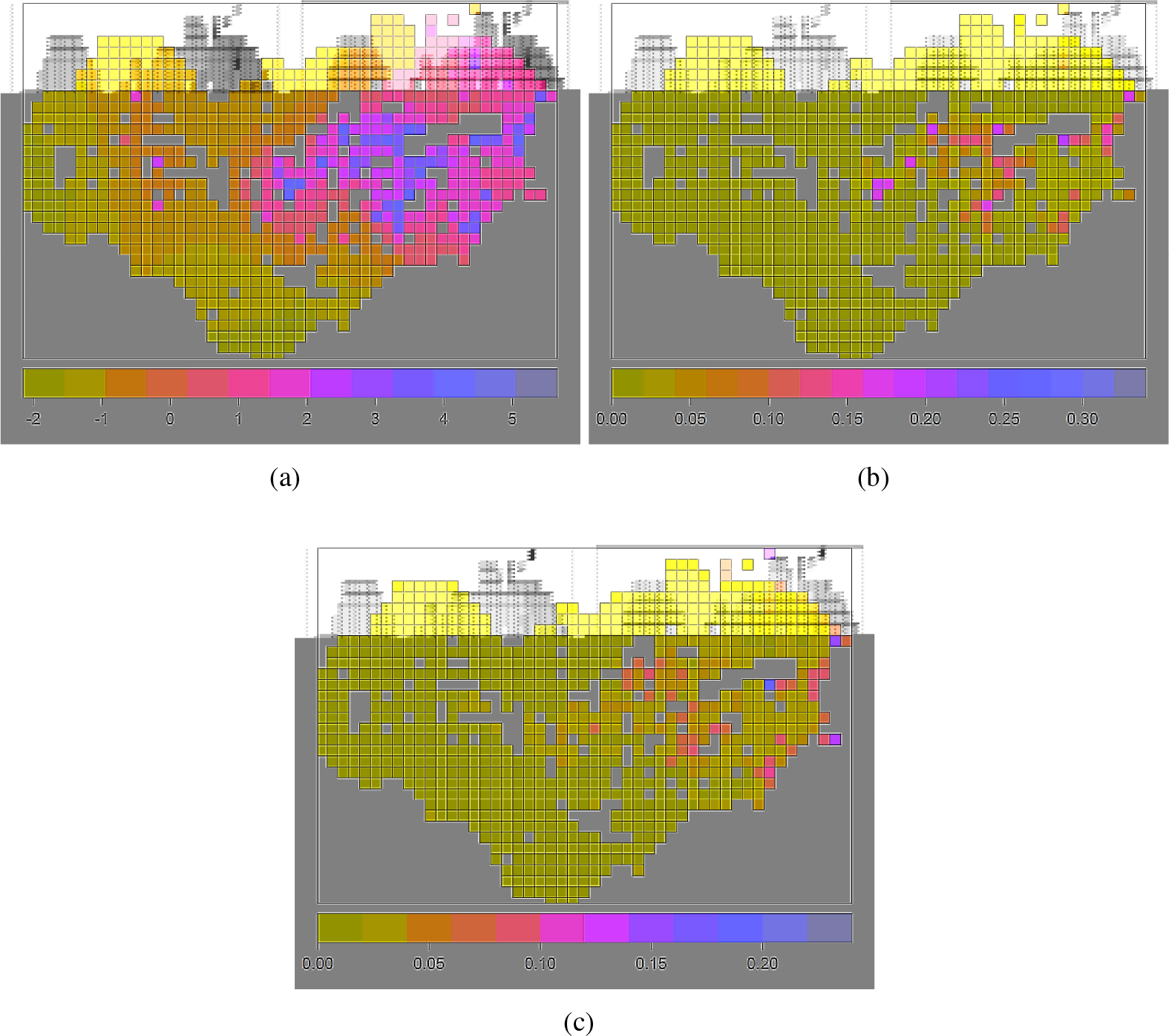
Geographical representation of the model of *Xylella fastidiosa* incidence distribution in the demarcated area in Alicante, Spain, with (a) the mean posterior distribution of the spatial effect, and (b) the mean posterior distribution of the incidence (0-1), and (c) its corresponding standard deviation.

Regarding *Xf* incidence estimates (Fig. 4(b) and 4(c)), mean posterior distribution varied from 0.002 to 0.215 and its corresponding standard deviation ranged from 0.003 to 0.191. The cells with the highest incidence values were concentrated in the central and eastern parts of the demarcated area, coinciding with those cells that presented the highest value for the spatial effect and proportion of positives. On the other hand, in the cells where *Xf* was not detected, mean incidence was estimated as zero or somewhat close to it, because in some cases the presence of positive neighboring cells contributed with the spatial dependence structure. The standard deviation showed higher values in those cells where the incidence estimate was more influenced by the spatial effect, that is, the cells that present higher values in Fig. 4(c), as well as in those having the highest incidence values.

#### Effect of sampling intensity on *Xf* incidence estimates

The effect of the sampling intensity on *Xf* incidence estimates (equation 5) was assessed by generating different data subsets from the reference database by limiting the sampling intensity with certain threshold values. These threshold values were established at 9 (*DS*_9_), 23 (*DS*_23_), 37 (*DS*_37_), and 51 (*DS*_51_) samples/cell. Note that 51 samples/cell corresponded to the optimum sampling intensity estimated for the 1 km^2^ grid resolution by the optimization algorithm (Section 2.2). The value of 9 samples/cell was consistent with the third quartile value of the sampling intensity of the reference database. The values of 23 and 37 samples/cell were chosen arbitrarily in the range of possible values between 9 and 51.

The different data subsets implied a reduction in the number of samples compared to the reference dataset. Furthermore, subsetting also implied a change in the distribution of both positive and negative samples and overall incidence, as shown in Supplementary material Table S2. Supplementary material Fig. S1 and S2 display the changes in the distribution of the sampling intensity (samples/cell), number of positive samples per cell, bacterium presence cells, and incidence per cell. Note that the quantities obtained with the different data subsets were summarized by the replicate showing the median behavior in relation to the total number of positive samples.

Table 7 shows the discrepancy measures used to assess the stability of the marginal posterior distributions of the model parameters and hyperparameters, including bias, standard error (SE), and standard deviation (SD), with their corresponding values associated to the different data subsets.

**Table 7:** Discrepancy measures bias, standard deviation (SD), and standard error (SE) for assessing parameters/hyperparameters marginal posterior stability with the data subsets (9, 23, 37 or 51 samples/cell) compared to the reference dataset of the demarcated area for *Xylella fastidiosa* in Alicante, Spain.

| Data subsets | Parameter/Hyperparameter | Bias | SE | SD |
| --- | --- | --- | --- | --- |
| $DS_9$ | $\beta_0$ | -0.364 | 0.476 | 0.318 |
| | $\sigma_v$ | -0.009 | 0.281 | 0.181 |
| $DS_{23}$ | $\beta_0$ | -0.128 | 0.360 | 0.185 |
| | $\sigma_v$ | -0.039 | 0.220 | 0.112 |
| $DS_{37}$ | $\beta_0$ | 0.008 | 0.324 | 0.008 |
| | $\sigma_v$ | -0.055 | 0.204 | 0.062 |
| $DS_{51}$ | $\beta_0$ | 0.017 | 0.319 | 0.056 |
| | $\sigma_v$ | -0.033 | 0.201 | 0.044 |

In relation to *β*_0_ marginal posterior stability, bias (absolute values) showed the highest values in the data subsets with more restrictive sampling intensities. On the other hand, *DS*_37_ and *DS*_51_ presented a similar behavior since both sampling intensities generated similar data subsets (Supplementary material Table S2). The SD and SE mimicked bias behavior. Regarding the stability of the hyperparameter posterior (σ_*v*_), the bias did not show a clear trend and all data subsets had similar values with a difference between the maximum (*DS*_37_) and minimum (*DS*_9_) of 0.046. The SE and SD also exhibited the highest trend associated with the data subsets that were more restrictive with sampling intensity.

Changes in the posterior distribution of *Xf* incidence and the spatial effect were assessed graphically (Fig. 5) by comparing the inference outcomes related to the replicas (averaged) in relation to the reference outputs (Fig. 4). In general, bias was greater in the data subsets that were more restrictive with sampling intensity although its range of values varied for each quantity evaluated. In general cells in the central and eastern parts of the demarcated area, with the highest sampling intensities (Supplementary material Fig. S1), exhibited higher bias related to the posterior distribution of the incidence (mean and SD).

**Fig. 5:**
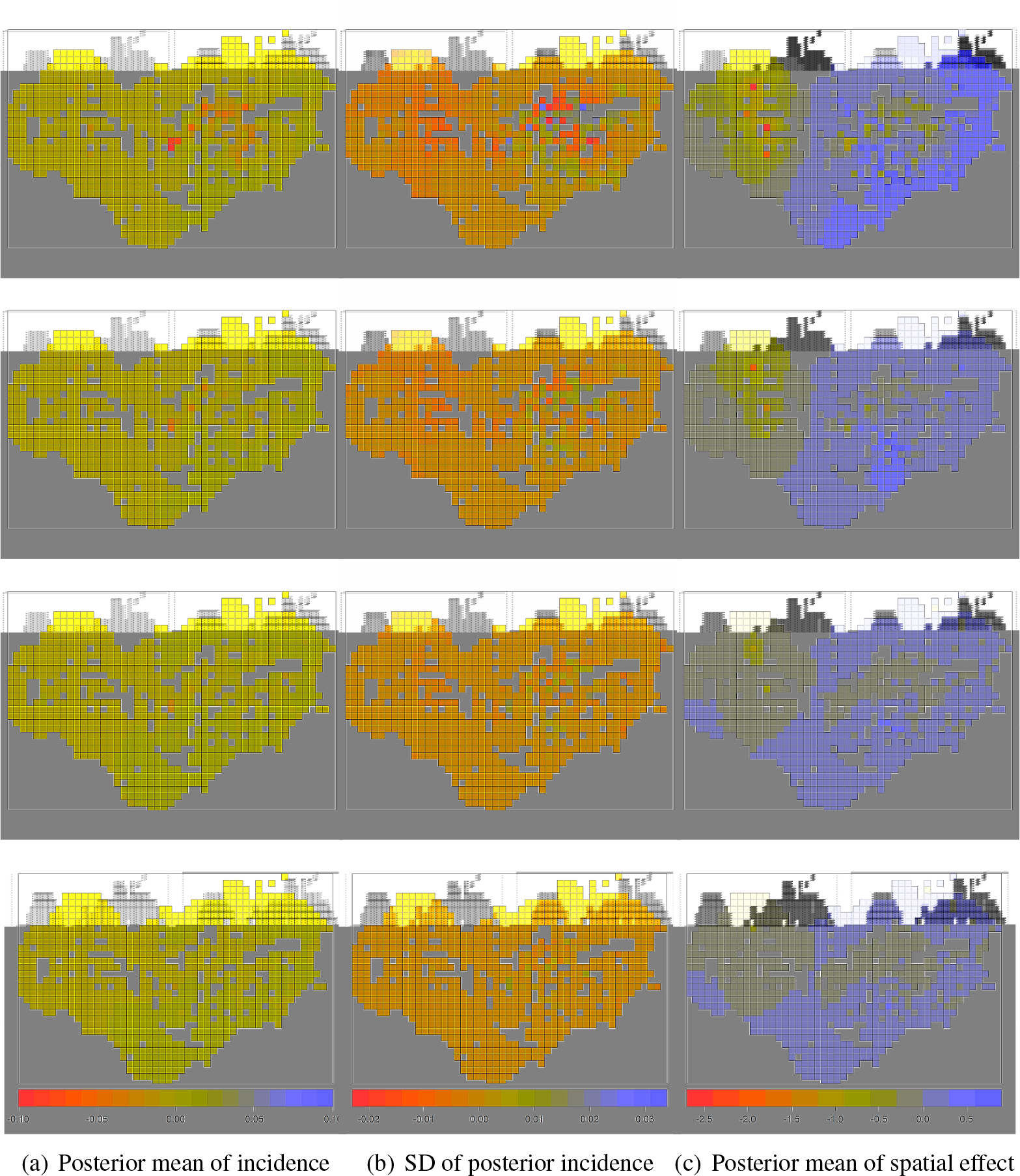
Bias for the posterior mean and standard deviation (SD) of incidence and posterior mean of the spatial effect for the data subsets *DS*_9_ (row 1), *DS*_23_ (row 2), *DS*_37_ (row 3), *DS*_51_ (row 4) relative to the model fitted to the reference dataset of the demarcated area for *Xylella fastidiosa* in Alicante, Spain.

With the data subsets *DS*_9_ and *DS*_23_, the stability of the spatial effect was negatively affected beyond the cells in the central and eastern parts of the demarcated area. Likewise, the data subsets *DS*_37_ and *DS*_51_ exhibited the most robust inferences with bias values close to zero and two clear groups of cells. One group was in the central and eastern parts of the demarcated area where the spatial effect was slightly underestimated. This group corresponded to cells with the highest sampling intensities and cells positive for *Xf*. The other group of cells in the north-west presented a slight overestimation of the spatial effect.

## 4 Discussion

Planning delimiting surveys implies reaching a compromise between the available resources and the extent of the area. After detecting an outbreak, the actual distribution of the disease is often unknown by plant health authorities, which further reduces the efficacy of control efforts, such as eradication or containment. Our generic delimiting survey strategy deals with these challenges and demonstrates that sequencing and adapting inspection and sampling to different spatial resolutions allows to be more accurate in delimitation infected zones given the typical aggregated spatial behavior of most plant diseases.

We developed a generic sequential adaptive strategy operationally deployed to delimit the geographical distribution of *Xf* in Alicante. The performance of two survey designs was evaluated and both improved the efficiency while maintaining the efficacy in relation to the current one. Our strategy involves sequencing inspection and sampling in time considering increasing spatial resolutions to define the grid resolution. Additionally, inspection and sampling intensities in finer spatial resolutions are deployed based on the information obtained at previous coarser resolutions. One important aspect of our work is that sampling intensity was defined by means of an optimization-based principle which maximizes *Xf* detection for each spatial resolution.

Our sequential adaptive strategy reduces inspection intensity while achieving the same spatial resolution indicated by the current legislation, owing to the phase-approach used to define grid cells at an increasing spatial resolutions. The current delimiting survey strategy implies carrying out parallel inspections in two predefined surveillance areas: the first 1^*st*^ km radius of the buffer zone using a grid resolution of 0.01 km^2^ cells and the outer buffer zone up to a radius of 5 km using a grid resolution of 1 km^2^. Conversely, our strategy defines an increasing spatial resolution in the whole demarcated area, from 1 km^2^ up to 0.01 km^2^, based on the information obtained in the previous inspection/sampling phase. That is, it allows delimitation of the spatial extent of the pathogen at coarser spatial resolutions while demarcating areas for implementing control measures at finer resolutions.

The sequential adaptive strategy resulted in a substantial reduction of the inspection intensities compared to the current one. This was particularly noted at the finest spatial resolution (0.01 km^2^), in which 2,225 and 7,100 cells were estimated to be inspected for the three-phase/two-phase designs, respectively, compared to the 17,700 cells of the current strategy. Overall, our sequential adaptive strategy will assist plant health authorities with a more efficient allocation of resources for disease control. Managers could deploy disease control tactics (plant removal, vector control, etc.) in a more targeted way. Likewise, the reduction in the survey effort would help to reduce delays in surveying, laboratory testing, and implementation of control measures while being more logistically feasible.

Our approach also allows the optimum sample size to be calculated for each survey resolution. The Decision prioritizes sampling plants with *Xf* -like symptoms, but no information is provided related to sample size calculation for delimiting surveys in the buffer zone. Thresholds of sampling intensity were estimated by a generic optimization algorithm which was sequentially adapted to each spatial resolution and with a stopping rule set to maximize the detection of infested cells. The algorithm uses random sampling because symptoms associated with *Xf* are not specific and can be confused with other biotic or abiotic disorders. Moreover, the incubation period (i.e., from infection to symptom expression) of *Xf* may be rather long and some plant species may not show symptoms even when infected by the pathogen (Purcell *et al*., 1999; Loconsole *et al*., 2016).

Optimum sampling intensity was estimated at 51, 45, 14 samples/cell for 1, 0.25 and 0.01 km^2^ cells. These results provide new insights to the ongoing work aimed to develop risk-based survey approaches for quarantine pest and pathogens (EU, 2019b) in which statistically based sample size calculations are being implemented (see EFSA, 2019, for further details). Once the delimitation of an outbreak has been accomplished, further surveys can provide additional information on the feasibility of management plans, disease dynamics in the infested area and the efficacy of targeted control tactics. Consequently, the optimum sampling intensity calculated for a resolution of 1 km^2^ was then validated by assessing its performance to estimate *Xf* incidence by means of a Bayesian hierarchical spatial model. The dual focus on applying advanced survey designs and modeling techniques was considered to maximize the quality of the information collected and the rigor of inference, as suggested by Johnson *et al*. (2013); Pacifici *et al*. (2016).

INLA has been proven to be a computationally efficient methodology to implement complex Bayesian hierarchical models which consider dependent structures among the data. Our model introduces spatial autocorrelation and also uses absence observations to estimate *Xf* distribution, thereby overcoming the deficiencies of other species distribution models (SDMs) used in that context which consider just presence-only data or generate pseudo-absence data and do not consider spatial dependence (Bosso *et al*., 2016; Gutiérrez-Hernández & García, 2019). Additionally, our modeling proposal also highlights the importance of adding model complexity to account for deficiencies in data collection. The integration of the spatial dependence structure was essential to mitigate the influence of heterogeneity in sampling intensities and also to explain the variability found in the distribution of *Xf*. Moreover, modeling outputs also evidenced the marginal influence of climatic covariates in that regard in the study area. Both aspects could be considered in future surveillance actions.

The evidence of the strong influence of the spatial effect means that the areas close to positive findings of *Xf* are more likely to be infested, illustrating the aggregated behavior of this pathogen. Nevertheless, here the spatial structure was included assuming a predefined distance of 2.5 km among cells. To define spatial correlation more accurately it could be interesting to consider additional information such as vector dispersal distances, although this information is still imprecise for the European outbreaks (EFSA, 2019). In contrast to other works (Godefroid *et al*., 2019), the scant relevance of climatic covariates found in our study prevents to establish a direct relationship with the distribution of *Xf* in the study area. Nevertheless, *Xf* subsp. *multiplex* is more widely distributed in the EU territory than *Xf* subsp. *fastidiosa* and *Xf* subsp. *pauca*, and it is known to have suitable climatic conditions in the vast majority of the territory (EFSA, 2019). As it may be the general situation in outbreak areas, the limited extent of the study area might also have played an important role in this respect.

In sum, our sequential adaptive survey strategy was able to efficiently delimit the extent of *Xf* and to estimate its incidence in the demarcated area in Alicante. This survey strategy can provide plant health authorities information about the local spatial variation of *Xf*. Given its sequential and adaptive nature, this strategy may assist to optimize survey resources and implement disease control and regulatory actions in a more targeted way. A more targeted allocation of management efforts could increase the efficiency and efficacy of control programs, such as eradication and containment, thus reducing treatment costs and minimizing side effects. Our strategy was designed to put more sampling effort in areas likely to be infested, but also allowing to adapt inspection and sampling intensity. Optimum values of sampling intensity could be used as a benchmark to be explored in other *Xf* outbreaks and compared with other inspection/sampling size calculation methods. Our work evidences the need to design feasible survey strategies which benefit from previous information and optimize the survey resources available.

## Acknowledgments

The present work has been funded by Horizon 2020 project No 727987 XF-ACTORS (*Xylella Fastidiosa* Active Containment Through a Multidisciplinary-Oriented Research Strategy) and the projects E-RTA 2017-00004-C06-01 FEDER INIA-AEI Ministerio de Economía y Competitividad and Organización Interprofesional del Aceite de Oliva Espa ñ ol, Spain. The work of ALQ and DC has been supported by grants MTM2016-77501-P and TEC2016-81900-REDT from the Spanish Ministry of Science, Innovation and Universities State Research Agency (jointly financed by the European Regional Development Fund, FEDER).

## Author contributions

EL, AV, ALQ, DC conceived the study. EL designed the analysis. VD and AFM provided the data. EL and MS analyzed the data. EL wrote the manuscript.

## Data and code availability

The data and the `R` scripts to reproduce the analyses are available at https://bitbucket.org/elaher/xylellafastidiosa_reproducibleresearch/src/master/

## Supplementary material

Additional Supplementary material may be found online in the Supplementary Material tab for this article.

**Table S1** Model comparison for *Xylella fastidiosa* incidence in the demarcated area in Alicante, Spain, based on WAIC and LCPO criteria.

**Table S2** Numerical comparison between data subsets and the reference database of the demarcated area for *Xylella fastidiosa* in Alicante, Spain, in terms of the total number of samples (% in relation to the reference database), positive samples, negative samples, and the global incidence.

**Fig. S1** Geographical distribution of sampling intensity and number of positive samples per cell for *SD*_9_ (row 1), *SD*_23_ (row 2), *SD*_37_ (row 3), *SD*_51_ (row 4), and reference situation (row 5).

**Fig. S2** Geographical distribution of bacterium presence and incidence per cell for *SD*_9_ (row 1), *SD*_23_ (row 2), *SD*_37_ (row 3), *SD*_51_ (row 4), and reference situation (row 5).

