## Supplementary material for "Tracking the outbreak. An optimized delimiting survey strategy for *Xylella fastidiosa*"

### Supporting information

#### Model selection

Table S1: Model comparison for *Xylella fastidiosa* incidence in the demarcated area in Alicante, Spain, based on WAIC and LCPO criteria.

| Model | WAIC | LCPO |
|---|---|---|
| $\beta_0 + bio1 + bio7 + v$ | 445.106 | 4.394 |
| $\beta_0 + bio1 + bio7 + v + u$ | 445.136 | 4.372 |
| $\beta_0 + bio1 + bio7 + bio13 + v + u$ | 445.190 | 4.403 |
| $\beta_0 + bio1 + bio7 + bio13 + v$ | 445.226 | 4.348 |
| $\beta_0 + bio7 + v + u$ | 445.497 | 4.415 |
| $\beta_0 + bio7 + v$ | 445.505 | 4.423 |
| $\beta_0 + bio7 + bio13 + v + u$ | 445.554 | 4.420 |
| $\beta_0 + bio7 + bio13 + v$ | 445.601 | 4.387 |
| $\beta_0 + bio1 + v + u$ | 445.641 | 4.546 |
| $\beta_0 + bio1 + v$ | 445.661 | 4.423 |
| $\beta_0 + bio1 + bio13 + v + u$ | 445.789 | 4.406 |
| $\beta_0 + bio1 + bio13 + v$ | 445.850 | 4.396 |
| $\beta_0 + v$ | 445.965 | 4.627 |
| $\beta_0 + bio13 + v + u$ | 446.003 | 4.396 |
| $\beta_0 + bio13 + v$ | 446.038 | 4.425 |
| $\beta_0 + v + u$ | 446.096 | 4.665 |

#### Sampling intensity vs. incidence modeling

Table S2: Numerical comparison between data subsets and the reference database of the demarcated area for *Xylella fastidiosa* in Alicante, Spain, in terms of the total number of samples (% in relation to the reference database), positive samples, negative samples, and the global incidence.

| **Data subsets** | **N samples (%)** | **N positives** | **N negatives** | **Global Incidence** |
|---|---|---|---|---|
| $\boldsymbol{DS}_9$ | 4432 (54%) | 65 | 4367 | 0.0146 |
| $\boldsymbol{DS}_{23}$ | 6123 (75%) | 126 | 5997 | 0.0206 |
| $\boldsymbol{DS}_{37}$ | 7002 (86%) | 168 | 6834 | 0.0240 |
| $\boldsymbol{DS}_{51}$ | 7519 (92%) | 196 | 7323 | 0.0260 |
| Reference database | 8146 (100%) | 237 | 7909 | 0.0291 |

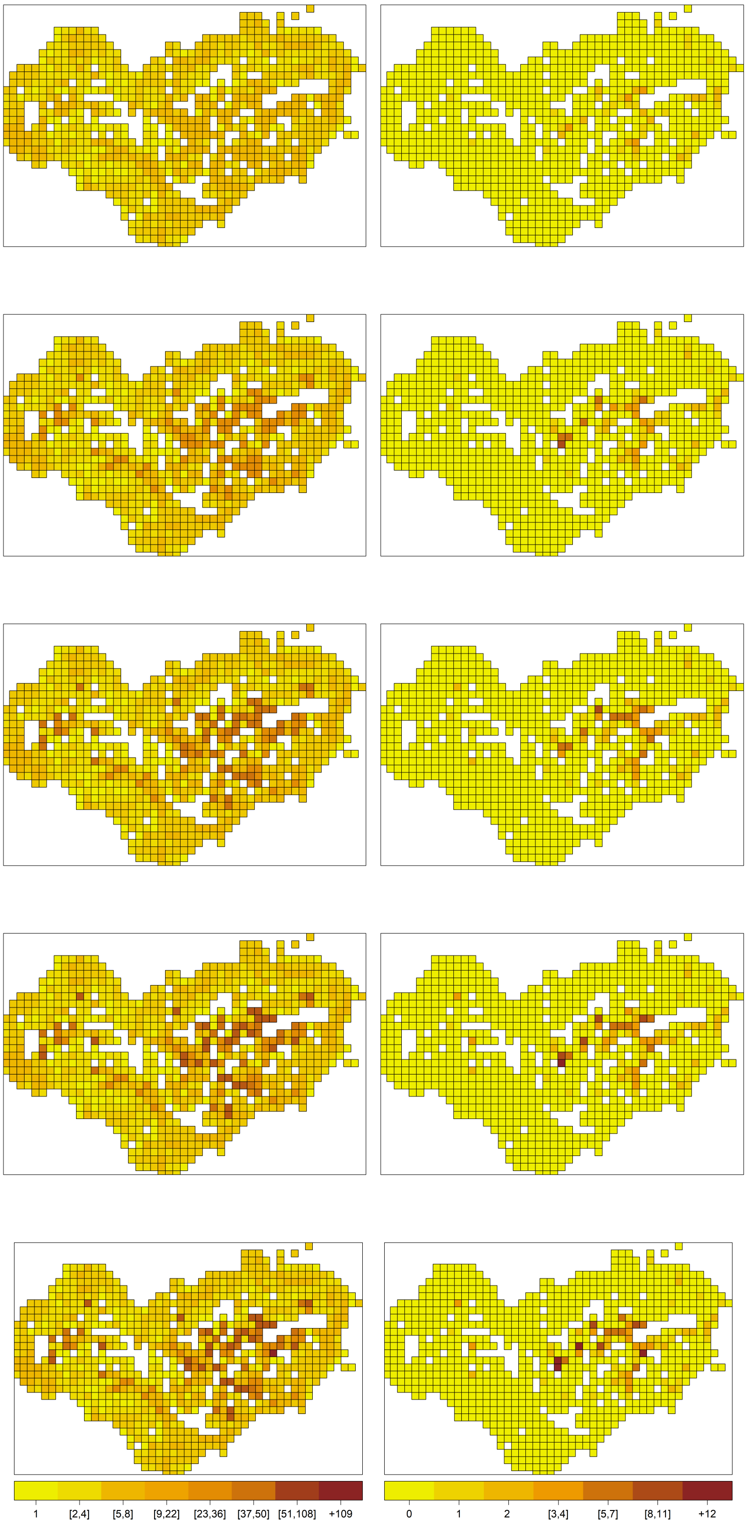




(a) Sampling intensity per cell (b) Number of positives samples per cell

Fig. S1: Geographical distribution of sampling intensity and number of positive samples per cell for $SD_9$ (row 1), $SD_{23}$ (row 2), $SD_{37}$ (row 3), $SD_{51}$ (row 4), and reference situation (row 5).

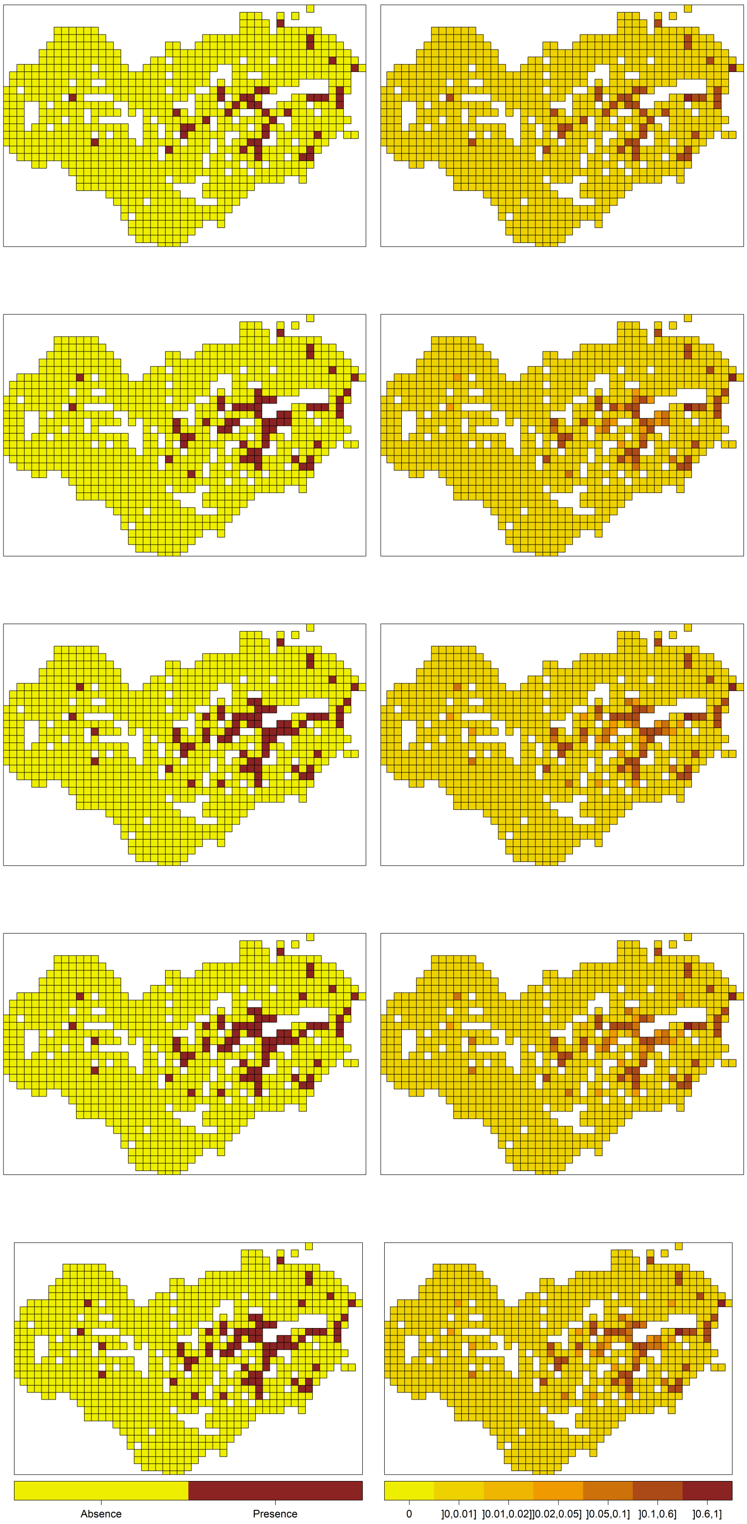


(a) Bacterium presence per cell (b) Incidence per cell

Fig. S2: Geographical distribution of bacterium presence and incidence per cell for $SD_9$ (row 1), $SD_{23}$ (row 2), $SD_{37}$ (row 3), $SD_{51}$ (row 4), and reference situation (row 5).
